## Supplemental information for "Single-cell Chromatin Accessibility and Lipid Profiling Reveals a Metabolic Shift in Adipocytes Induced by Bariatric Surgery"

### Supplementary Text

#### Freezing nuclei after tagmentation

Batch effects are a significant concern in single-cell sequencing experiments and can arise when nuclei extractions, tagmentation, library amplifications are spread across multiple days. We assessed a protocol alteration to minimize batch effects by consolidating library preparation steps in fewer days. We cryopreserved nuclei after tagmentation to allow samples to be prepared in tandem. Though cryopreservation of cells for bulk ATAC-seq did not lead to reduced quality data, this has not yet been tested for single-cell ATAC-seq (Milani et al., 2016). Notably, compared to freshly isolated nuclei, frozen nuclei showed no discernable differences in quality with a high degree of correlation in counts per gene (fig S1e). We applied this strategy to our adipose snATAC-seq library preparations, increasing the per day throughput of each library preparation step.

**Supplementary Figures**

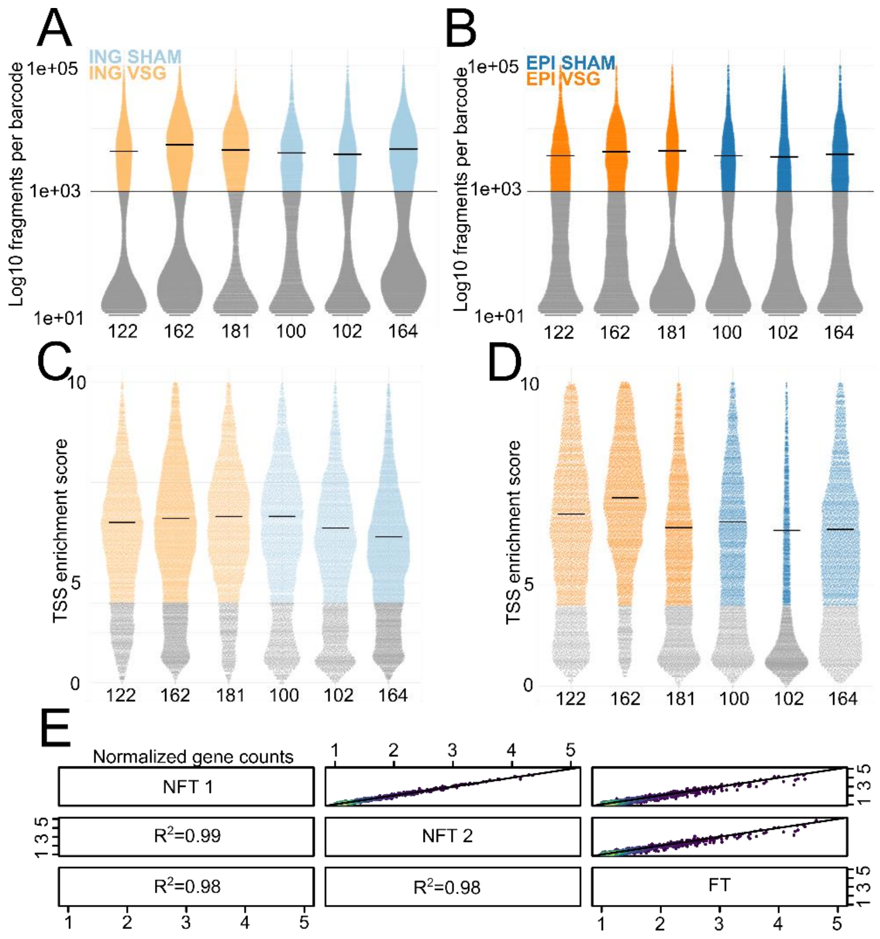


**Supplementary Figure 1. Quality control metrics for each biological replicate**

(A,B) Fragment count distributions per barcode for each cell for each biological replicate shown in Fig 1 in (A) ING and (B) EPI. The median value for each group is shown with a thick bar.

(C, D) TSS enrichment score per barcode for each cell for each biological replicate shown in Fig 1 in (C) ING and (D) EPI. Numbers below violin plots denote the individual mouse identification number.

(E) Spearman correlation of normalized gene counts comparing non-freeze thaw technical replicate libraries (NF1 and NF2) vs. the freeze-thaw library (FT) from the same pool of tagmented nuclei.

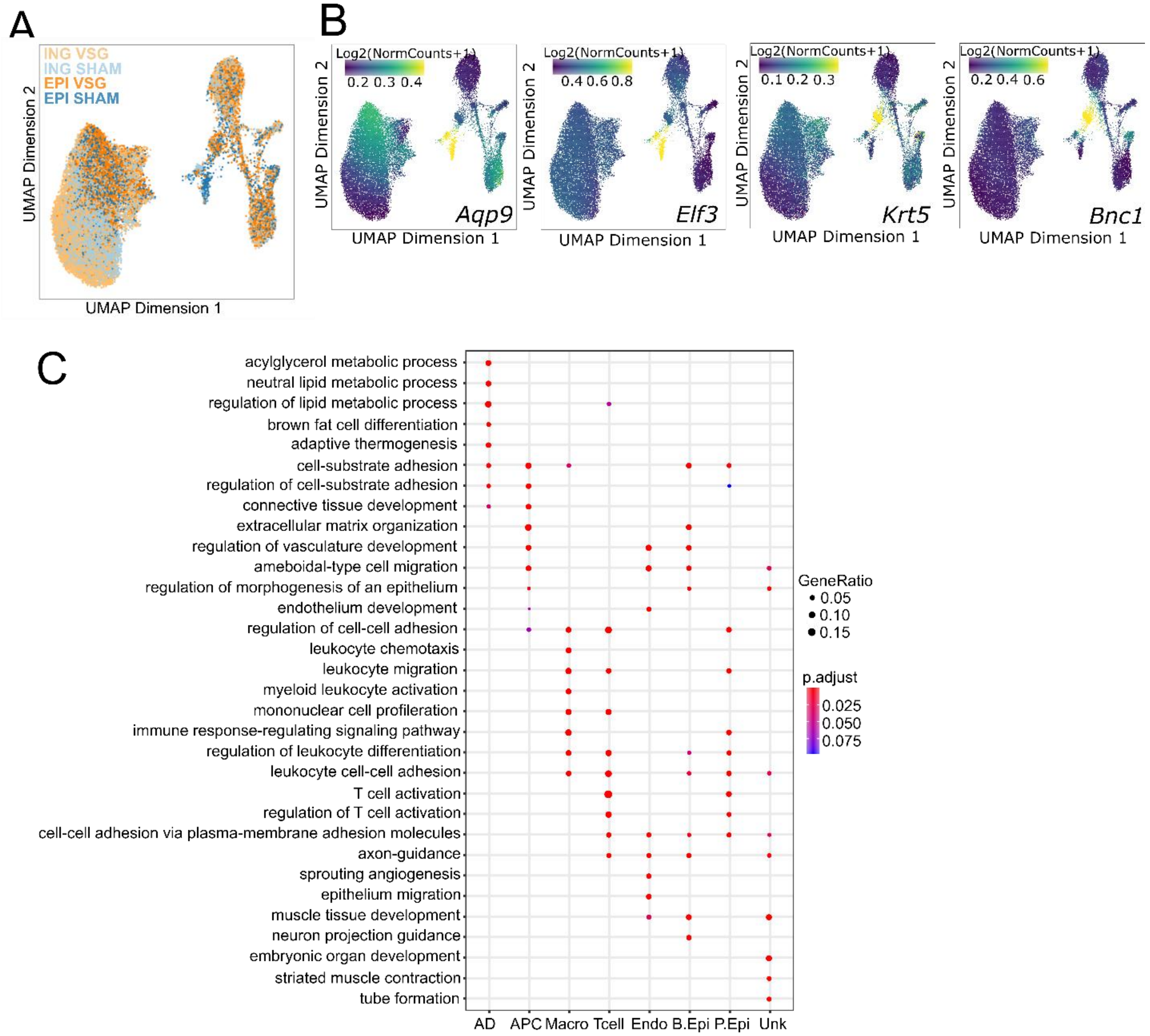


**Supplementary Figure 2.**

**(**A) UMAP plot showing distribution of cells from inguinal (light blue, ING-SHAM; light orange, ING-VSG) and epididymal (dark blue, EPI-SHAM; dark orange, EPI-VSG).
(B) UMAP with imputed accessibility of basal epididymal (*Aqp9* and *Bnc1)* and principal epididymal (*Krt5* and *Elf3)* cell type markers.
(C) GO enrichment for top 500 DA genes from each major cluster.

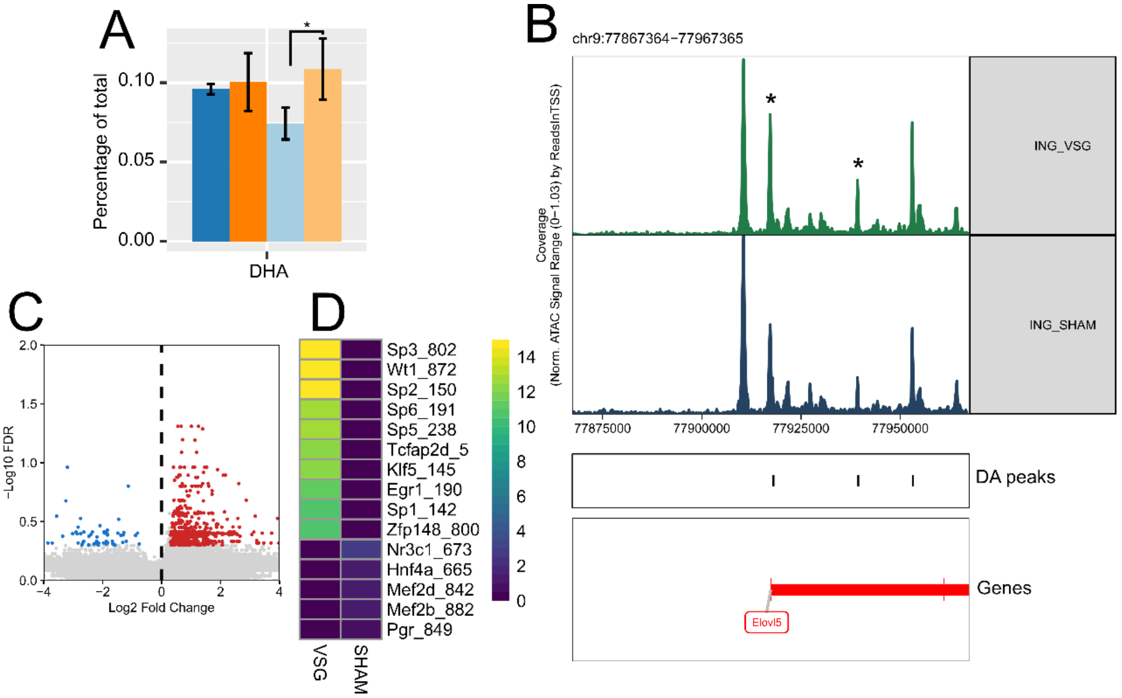


**Supplementary Figure 3. Differential accessibility of transcription factor motifs and changes in DHA fatty acid in response to VSG**

(A) Percent of DHA fatty acid in adipose tissues.

(B) Genome browser shot of the promoter region of *Elovl5* showing STAT5A motifs (denoted by *) within peaks that are more accessible in VSG. Differentially accessible, DA.

(C) Differential accessible peaks in EPI adipocytes. There were 593 peaks more accessible in EPI VSG (positive fold change) and 66 peaks more accessible in EPI SHAM (negative fold change)

(D) Enriched transcription factor motifs in differentially accessible peaks in EPI adipocytes.

Differentially accessible, DA; DHA, Docosahexaenoic acid

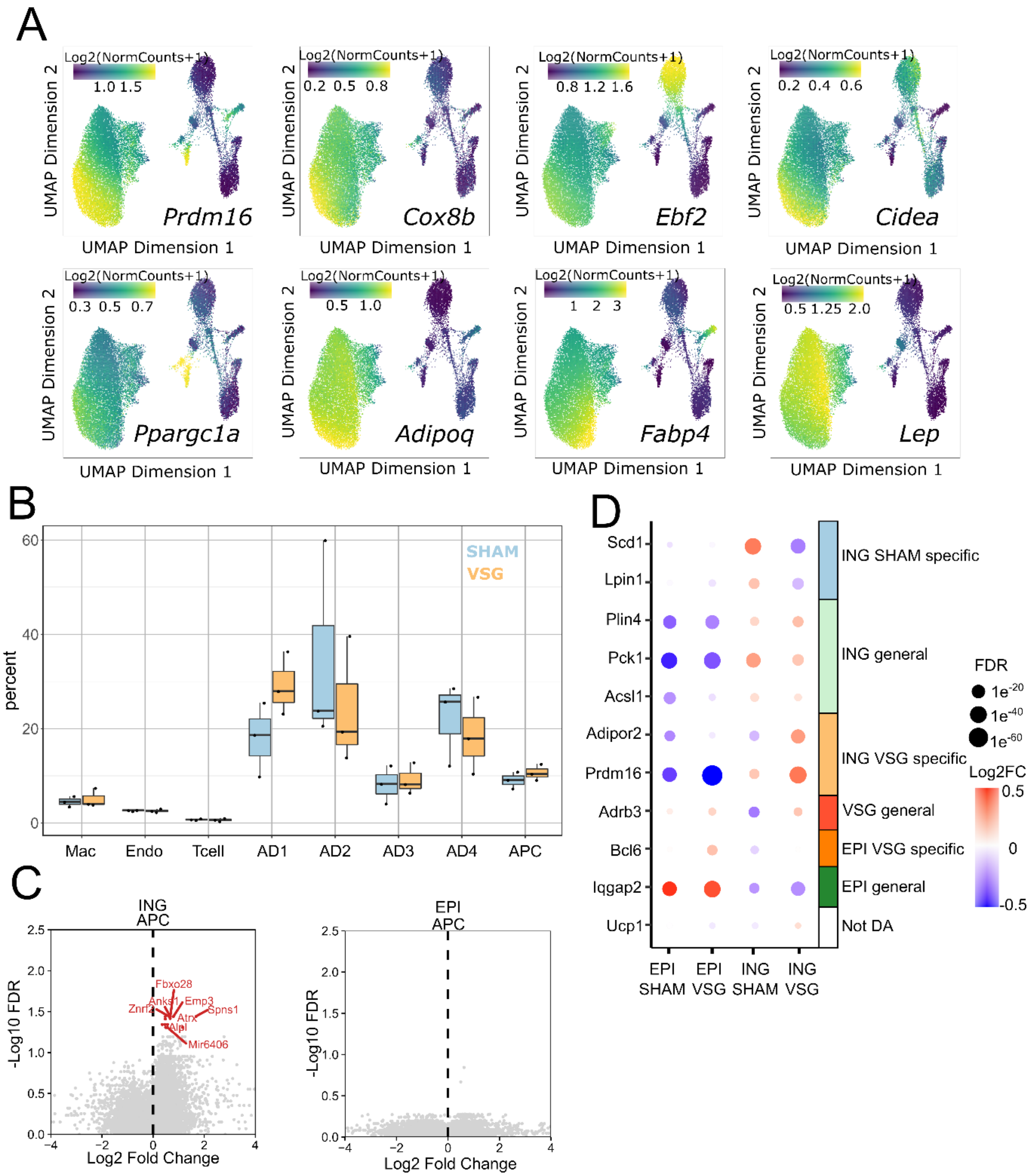


**Supplementary Figure 4. Differential accessibility of genes response to VSG in the ING depot**

(A) Imputed marker gene accessibility using MAGIC overlaid on the UMAP plot for beige-associated adipocyte markers (*Prdm16, Cox8b, Ebf2, Cidea, Ppargc1a*) and white-associated adipocyte markers (*Adipoq, Fabp4, Lep*).

(B) Relative distribution of cell types in the ING depot from SHAM and VSG animals. Percentages were normalized to the total number of cells within each biological replicate.

(C) Gene accessibility differences between VSG (positive fold change) and SHAM (negative fold change) in adipocyte progenitor cells (APC) of ING (left) and EPI (right).

(D) Relative average accessibility of genes with depot specific responses to VSG.
